# Convective forces contribute to post-traumatic degeneration after spinal cord injury

**DOI:** 10.1101/2024.06.17.599340

**Authors:** Hoi Y Kwon, Christopher Streilein, R. Chase Cornelison

**Affiliations:** Department of Chemical Engineering, University of Massachusetts Amherst, Amherst, MA 01003; Department of Biomedical Engineering, University of Massachusetts Amherst, Amherst, MA 01003

**Keywords:** Spinal cord injury, secondary neural injury, interstitial pressure, interstitial fluid flow, computational fluid dynamics, fluid shear stress, convection enhanced delivery

## Abstract

Spinal cord injury (SCI) initiates a complex cascade of chemical and biophysical phenomena that result in tissue swelling, progressive neural degeneration, and formation of a fluid-filled cavity. Previous studies show fluid pressure above the spinal cord (supraspinal) is elevated for at least three days after injury and contributes to a phase of damage called secondary injury. Currently, it is unknown how fluid forces within the spinal cord itself (interstitial) are affected by SCI and if they contribute to secondary injury. We find spinal interstitial pressure increases from -3 mmHg in the naive cord to a peak of 13 mmHg at 3 days post-injury (DPI) but relatively normalizes to 2 mmHg by 7 DPI. A computational fluid dynamics model predicts interstitial flow velocities up to 0.9 μm/s at 3 DPI, returning to near baseline by 7 DPI. By quantifying vascular leakage of Evans Blue dye after a cervical hemi-contusion in rats, we confirm an increase in dye infiltration at 3 DPI compared to 7 DPI, suggestive of higher fluid velocities at the time of peak fluid pressure. *In vivo* expression of the apoptosis marker caspase-3 is strongly correlated with regions of interstitial flow at 3 DPI, and exogenously enhancing interstitial flow exacerbates tissue damage. *In vitro*, we show overnight exposure of neuronal cells to low pathological shear stress (0.1 dynes/cm^2^) significantly reduces cell count and neurite length. Collectively, these results indicate that interstitial fluid flow and shear stress may play a detrimental role in post-traumatic neural degeneration.

**Translational Impact Statement:** Trauma to the central nervous system induces neural tissue degeneration, resulting in permanent disability and loss of function. A better understanding of this degenerative process is needed, towards developing new clinical treatments that effectively minimize tissue damage and preserve neural function after injury. The present study identifies a potential role for altered fluid transport within the injured spinal cord. These results provide new insight into basic pathophysiology and may inform therapeutic development for neuroprotection.

## Introduction

The National Spinal Cord Injury Statistical Center estimates that 18,000 people sustain a spinal cord injury (SCI) in the United States each year.^1^ Over a million Americans currently live with paralysis due to SCI,^1,2^ with deficits ranging from loss of motor function, difficulty breathing, and bladder and/or sexual dysfunction. Unfortunately, a viable treatment option to improve the quality of life of these patients is yet to be achieved.^3–5^

One challenge to treating SCI is the occurrence of secondary injury – a cascade of physical and biochemical phenomena that cause additional cell death and expansion of the tissue lesion over several weeks.^5–8^ In the acute and subacute phases of injury, disruption of the blood-spinal cord barrier and influx of immune cells leads to an increase in water content, or edema, and swelling of the tissue.^9,10^. The extent of edema correlates with injury severity and the degree of functional losses across animal models and humans.^9–12^

Edema occurs above the tissue but also within the interstitial space of the tissue, or region between the cells. The interstitial fluid is a vital bodily fluid that connects blood, cerebrospinal fluid, lymph fluid, and intracellular fluid.^13,14^ An important function of interstitial fluid is to maintain effective functioning of cells by enabling delivery of nutrients and removal of cell waste products.^13–15^ In a normal tissue environment, the flow of interstitial fluid informs cell physiology and function through shear stresses exerted at the cell surface.^14–16^ Similarly, pathological rates and patterns of interstitial flow can trigger inflammation.^17–19^ For example, disturbed flow in blood vessels promotes pro-inflammatory signaling in endothelial cells,^18^ and tumor-associated increases in interstitial flow causes pro-tumor transformation of local fibroblasts.^19^ While well-studied in a cancer environment, very little is currently known about how interstitial fluid flow affects pathology after SCI.^20–24^ The increase in fluid pressure after SCI due to blood-spinal cord disruption has potential to increase interstitial flow into the adjacent healthy tissue and promote pathological responses.

In this study, we examine the hypothesis that increases in fluid pressure after SCI lead to increases in interstitial fluid flow, which may contribute to tissue damage during the secondary injury phase. We present *in silico* and *in vivo* methods to understand how interstitial pressure influences interstitial flow in the spinal cord after injury. We also assess the role of increased fluid flow and shear stress on cell damage and lesion expansion after SCI using *in vitro* cell cultures and an *in vivo* technique called convection enhanced delivery. Our results ultimately improve our knowledge of interstitial flow in the context of SCI and the potential role it plays in secondary neural injury.

## Results

### Interstitial fluid pressure increases after SCI, peaking at 3 days post-injury

Fluid flow is governed by a pressure gradient, so we first wanted to determine how interstitial fluid pressure (ISP) in the spinal cord is affected by injury. Previous studies have reported values for supraspinal or intrathecal pressure (above the cord) after injury, but ISP has not been measured. We measured ISP in spinal cord tissue using a catheter-based device. The modes of measured ISP at different time points are plotted (**Figure 1B**), with the bar representing an average of the mode values for all recordings/animals per group. The average pressure of the normal tissue is -3 mmHg, and it significantly increases to 7 mmHg by 1-hour post injury (HPI) (p<0.01). The pressure peaks at 13 mmHg by 3 days post-injury (DPI), and drops to 2 mmHg by 7 DPI. The trend of pressure increasing by 1 HPI, remaining elevated until 3 DPI, and decreasing by 7 DPI is similar to that previously reported for intrathecal pressure measurements.^25^ There was a significant difference of ISP between normal and 1 HPI (p<0.01), 1 HPI and 3 DPI (p<0.05), and 3 DPI and 7 DPI (p<0.01). The greatest difference was between normal pressure and 3 DPI (p<0.0001). ISP was no longer statistically different from pre-injury values by 7 DPI (p=0.12).

**Figure 1:**
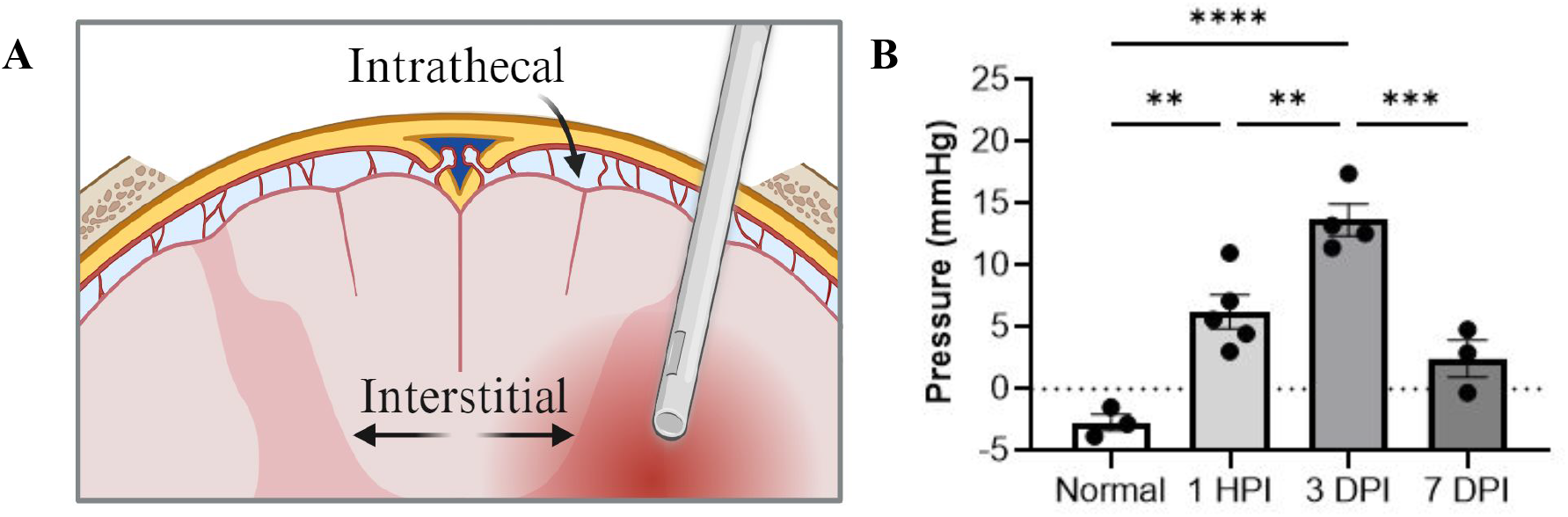
Interstitial fluid pressure increases after contusion spinal cord injury. A) Diagram of catheter-based measurement of interstitial fluid pressure in the spinal cord after laminectomy. Regions of intrathecal vs. interstitial pressure are labeled, and the catheter is shown inserted into the injury. Graphic made using an institutional license for Biorender. B) Interstitial fluid pressures in the naive (uninjured) and injured rat spinal cord in mmHg. Data were compared by ordinary one-way ANOVA and Tukey’s multiple comparisons test with a single pooled variance. **p<0.01, ***p<0.001, ****p<0.00001.

### Computational modeling predicts high interstitial fluid velocity and shear stress after SCI

Using the measured pressure values and parameters from previous literature,^26–29^ we constructed a computer model to simulate interstitial fluid flow velocity at 3 and 7 DPI. The outward pressure of the cavity was set at 1800 Pascal (∼13 mmHg) for 3 DPI and 600 Pascal (∼4.5 mmHg) for the 7 DPI model. The model was simulated using Brinkman’s equation (**Equation 1**) and the tissue parameters in **Table 1**. Representative results are shown in **Figure 2** below.

**Figure 2:**
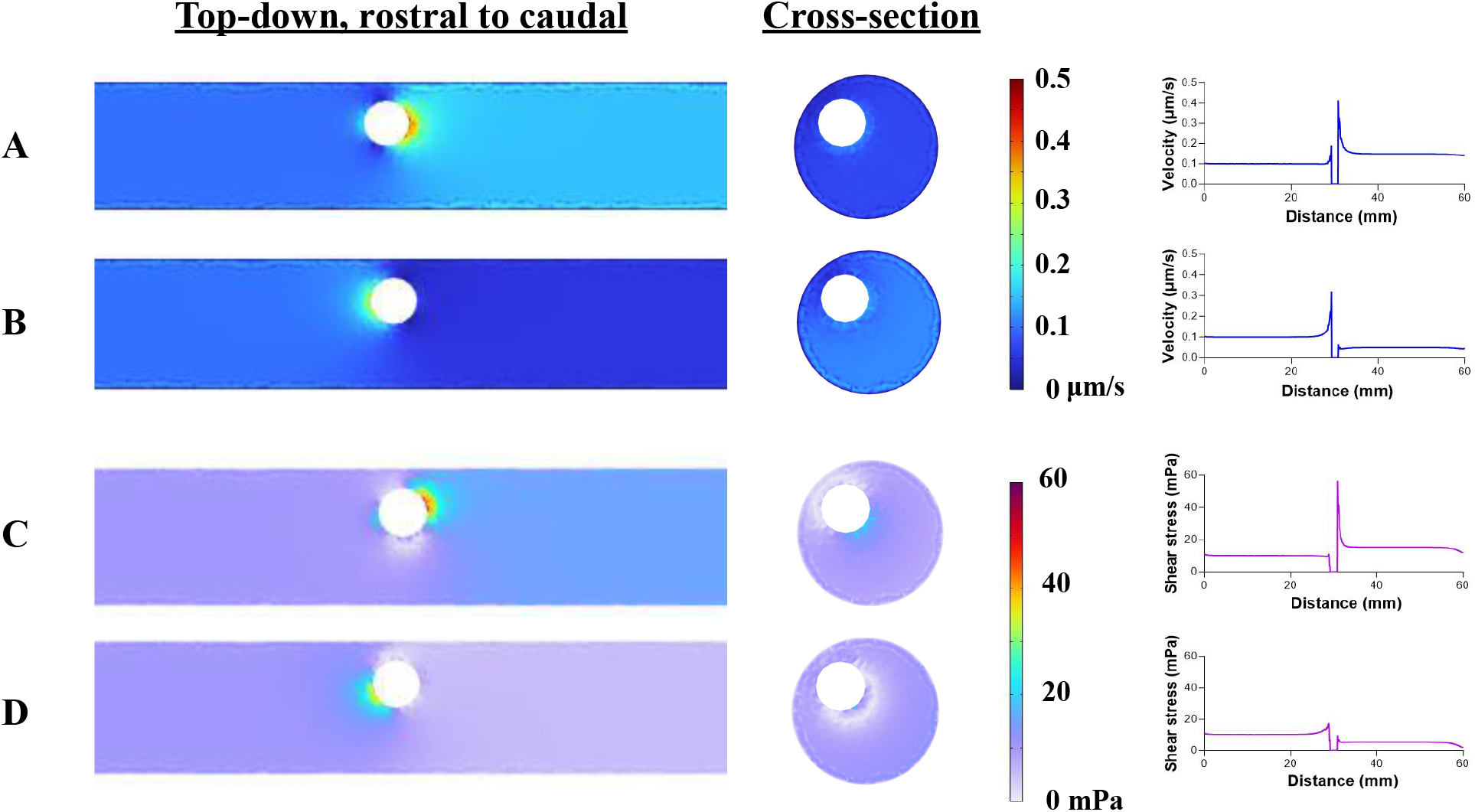
High fluid pressure after spinal cord injury is predicted to increase interstitial fluid flow and shear stress adjacent to the injury. Visual and graphical representations of simulated fluid forces after rat spinal cord injury for A) Fluid velocity at 3 days post-injury (DPI), B) Fluid velocity at 7 DPI, C) Fluid shear stress at 3 DPI, and D) Fluid shear stress at 7 DPI. COMSOL simulation results are shown from a top-down view, rostral (left) to caudal (right), and cross-sectional view. Corresponding graphs show representative values along the longitudinal axis from a single trial simulation.

**Table 1:** Transport parameters used to develop an *in silico* model of the rat spinal cord.

| Tissue parameter | Reported value | Reference |
| --- | --- | --- |
| Density, white matter ($\rho_{\text{wm}}$) | 1050 kg/m<sup>3</sup> | [26] |
| Porosity ($\varepsilon_p$) | 0.26 | [27] |
| Permeability, white matter ($k_{\text{wm}}$) | $2.5 \cdot 10^{-15} \text{ m}^2$ | [27,28,29] |
| Dynamic viscosity, fluid ($\eta_{\text{fl}}$) | 0.001 Pa*s | [28] |
| Normal interstitial pressure | -360 Pa |  |
| 3 DPI interstitial pressure | 1800 Pa |  |
| 7 DPI interstitial pressure | 300 Pa |  |

We first modeled fluid flow in a static cord, without a defined inlet rate of interstitial flow, to focus on understanding how pressure alone affected fluid flow (**Supplemental Figure 1**). To understand how physiological interstitial flow affects fluid velocity around the cavity, we then added a normal rate of interstitial flow at the inlet x=0 (rostral) in the x-direction. An inlet flow of 0.1 μm/s created anisotropic flow out of the cavity, biased in the caudal (+x) direction. In the 3 DPI model (**Figure 2A, C**), interstitial velocity around the cavity reaches a peak around 0.45 μm/s and plateaus near 0.15 μm/s downstream from the cavity. In terms of flow rate, the peak is 0.763 μL/min, and the plateau is 0.254 μL/min. The peak is more than 2 times that of a normal physiological interstitial flow rate (0.33 μL/min), whereas the flow rate at the plateau is comparable to the normal rate. For the 7 DPI model (**Figure 2B, D**), the peak velocity is around 0.3 μm/s (0.509 μL/min), or 1.7 times the normal flow rate, but quickly drops to a velocity of 0.1μm/s that is lower than the initial inlet velocity.

To understand how an increase in flow influences the forces sensed by cells, we used COMSOL to estimate the fluid shear stresses within the tissue. Shear stress in a normal physiological context is estimated to be around 0.01 dynes/cm^2^. As shown in **Figure 2C-D**, the highest peak at 3 DPI is around 60 mPa, equivalent to 0.6 dynes/cm^2^. This value is six times higher than what is as low pathological shear stress (0.1 dynes/cm^2^) and comparable to estimates for high physiological shear stress (0.6 dynes/cm^2^).^22,30,31^ The shear stress quickly dissipates to a minimum moving outward from the cavity, though it remains elevated in the 3 DPI model compared to physiological values. Shear stress close to the cavity in the 7 DPI setting reaches approximately 20 mPa, slightly higher than the low pathological shear stresses. While these values quickly return to physiological levels away from the cavity, cells near the cavity may still experience higher than normal fluid shear stresses, even at 7 DPI.

### Evans Blue infiltrates farther into the spinal cord interstitium at 3 DPI than 7 DPI

To begin assessing interstitial flow after injury *in vivo*, we used the dye Evans Blue, which binds to albumin in the bloodstream and extravasates into the interstitial fluid in regions of vascular leakage. The resulting extent of tissue staining therefore provides an approximation of interstitial flow out of the cavity and into the adjacent healthy tissue. **Figure 3** shows fluorescence images of injured spinal cord tissue at 3 DPI (**3A**) and 7 DPI (**3B**) with Evans Blue (magenta) and stained for neurofilament (green). At 3 DPI, the lesion area is identified by a lack of neurofilament staining, showing the absence of axons and neuronal soma. Evans Blue can be found outside of the lesion area, indicating that there is flow out of the cavity and into the surrounding tissue. The injured area of 7 DPI can be spotted with the neurofilament marker as it shows two different types of striations that differentiate between normal and lesion area. The aligned, straight structure are the axons and the uneven, fragmented filaments are likely the injured neural cells within the lesion. Evans Blue at 7 DPI is mostly contained inside the lesion area, with only some dye leaving the cavity. Hence, there is less flow out of the lesion at 7 DPI. We measured the distance Evans Blue traveled perpendicular to the lesion area as an indirect estimate of fluid flow. Shown in **Figure 3C**, the mean distance of Evans Blue extravasation averaged across four animals was 2129 μm at 3 DPI and 750 μm at 7 DPI, representing a significant statistical difference (p<0.01). This result aligns with our results from the *in silico* model and suggests increased fluid flow out of the lesion into the tissue at 3 days than at 7 days.

**Figure 3:**
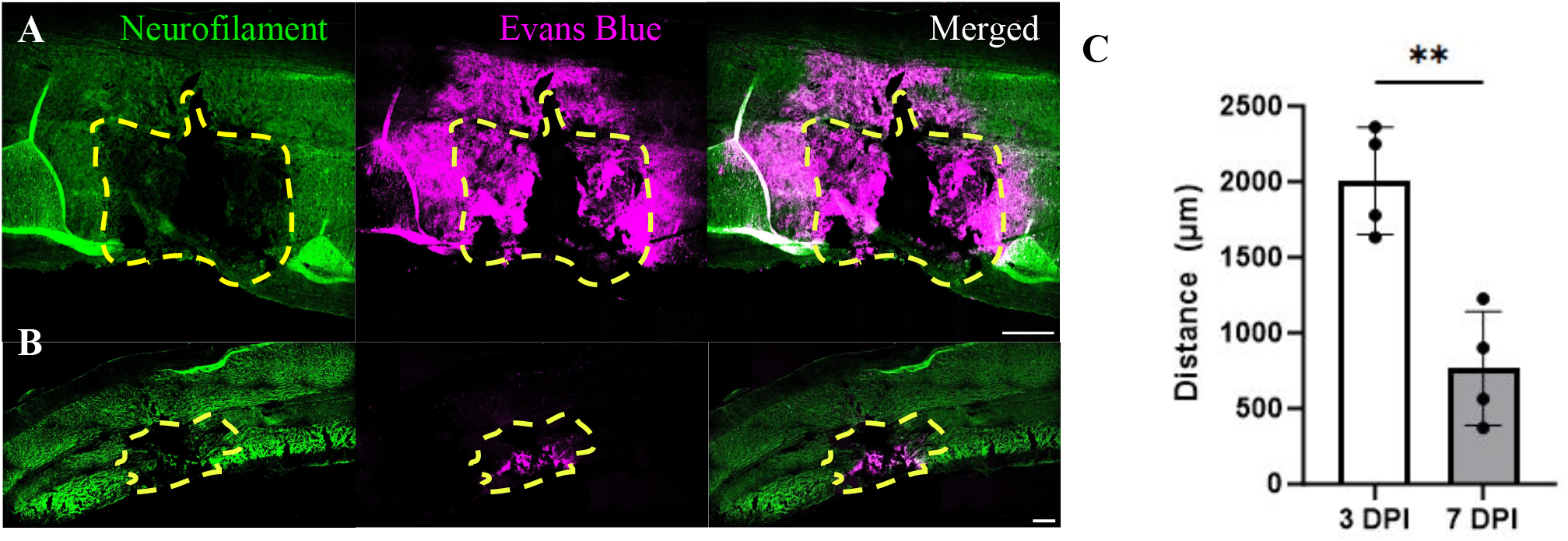
Evans blue extravasates farther into injury-adjacent tissue at 3 days compared to 7 days post-injury. A,B)The staining of spinal cord injury at 3 DPI (A) and 7 DPI (B) with neurofilament A and Evans Blue. The yellow dotted line represents the predicted lesion area. The Evans Blue spreads further in 3 DPI tissue than 7 DPI signifying the higher fluid flow out of the injury 24 hour prior to the sectioning. The white bar represents 200 **μ**m. C) The measurement of the distance Evans Blue traveled outside of the lesion area is plotted on the graph. At 3 DPI, the distance averaged at 2129 **μ**m and 750 **μ**m at 7 DPI. **p < 0.01

### Apoptosis during the sub-acute phase strongly correlates with regions with flow

To determine if an increase in flow around the cavity correlates with cellular damage, we stained the tissue sections for cleaved caspase-3, a marker of apoptosis. As expected, we observed widespread expression of cleaved caspase-3 around the cavity as an indication of active cell death and secondary injury processes (**Figure 4A**). We measured the degree of colocalization between caspase-3 and Evans Blue using Nikon imaging software to determine if the regions experiencing apoptotic cell death correlate with regions of interstitial flow. As shown in **Figure 4B**, the average Pearson’s correlation coefficient for caspase-3 versus Evans Blue was 0.60 at 3 DPI and 0.50 at 7 DPI, indicating a moderate to strong colocalization. We also calculated the Mander’s overlap coefficient (MOC), which showed similar results. The MOC values for caspase-3 and Evans Blue were 0.73 at 3 DPI and 0.76 for 7 DPI **(Figure 4C**), indicating a strong correlation. Therefore, the tissue regions containing active cell death exhibit moderate to strong overlap with tissue regions experiencing post-traumatic interstitial flow. Additionally, there was no significant difference between the correlation values at 3 and 7 DPI, so this colocalization is maintained throughout the relevant time frame for secondary neural injury.

**Figure 4:**
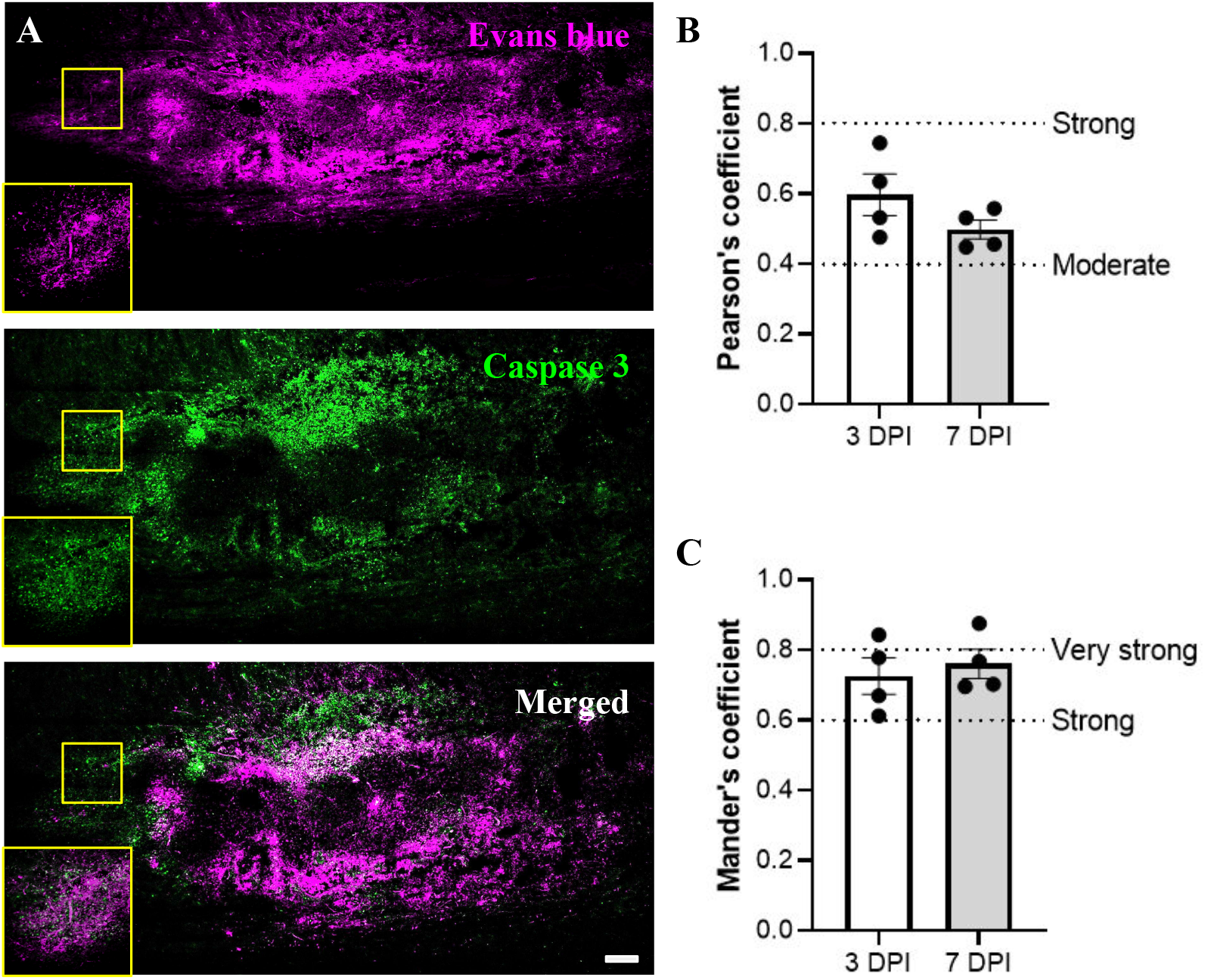
Perilesional interstitial flow strongly correlates with expression of cleaved caspase 3, a marker of apoptosis. A) Representative fluorescence images of 3 DPI spinal cord tissue labeled with Evans blue, indicating flow regions, and stained for cleaved caspase 3, a marker of apoptotic cells. Inset images show nuclear localization of caspase 3 at higher magnification. Scale bar is 200 μm. B-C) Correlational analyses were conducted to examine overlap between Evans blue and caspase staining using Pearson’s coefficient (B) and Mander’s coefficient (C). Value ranges indicative of Moderate, Strong, and Very Strong correlations are labeled.

### Artificially increasing flow induces lesion volume expansion

The above results demonstrate a correlation between apoptosis and fluid flow, but we also wanted to address the potential causation of heightened fluid flow and post-traumatic cell death. We therefore implemented a clinical technique called convection enhanced delivery, or CED, to externally facilitate an increase of interstitial flow, then we examined the consequence of this heightened flow on tissue damage.^32–34^ CED uses a catheter to deliver a fluid infusion at a defined rate into a tissue and drive increased interstitial flow. A cohort of animals was separated into two groups, each receiving a second surgery seven days after the initial injury, when the increase in interstitial pressure has mostly subsided. The control group received a catheter insertion only with no fluid delivery, while the +Flow group received saline infusion at 1.0 μL/min, or three times the normal rate of interstitial flow. The tissues were harvested three days after CED and stained with Luxol Blue to mark the demyelinated lesion area. As shown in **Figure 5**, the spinal lesion in the +Flow group extends further in the caudal and medio-lateral dimensions compared to untreated controls. We used the Cavalieri method (i.e., stereology) to calculate the mean lesion volume for both groups. The +Flow group reached 41.1 mm^3^ and was significantly larger than the 9.91 mm^3^ volume measured in the control group (p<0.01). In other words, increasing fluid flow within the injured spinal cord exacerbates tissue damage and promotes lesion volume expansion. Collectively, our results demonstrate that an increase in fluid flow, as predicted *in-silico* and visualized *in vivo*, both correlates with and can contribute to subacute tissue damage in a rat model of contusion SCI.

**Figure 5:**
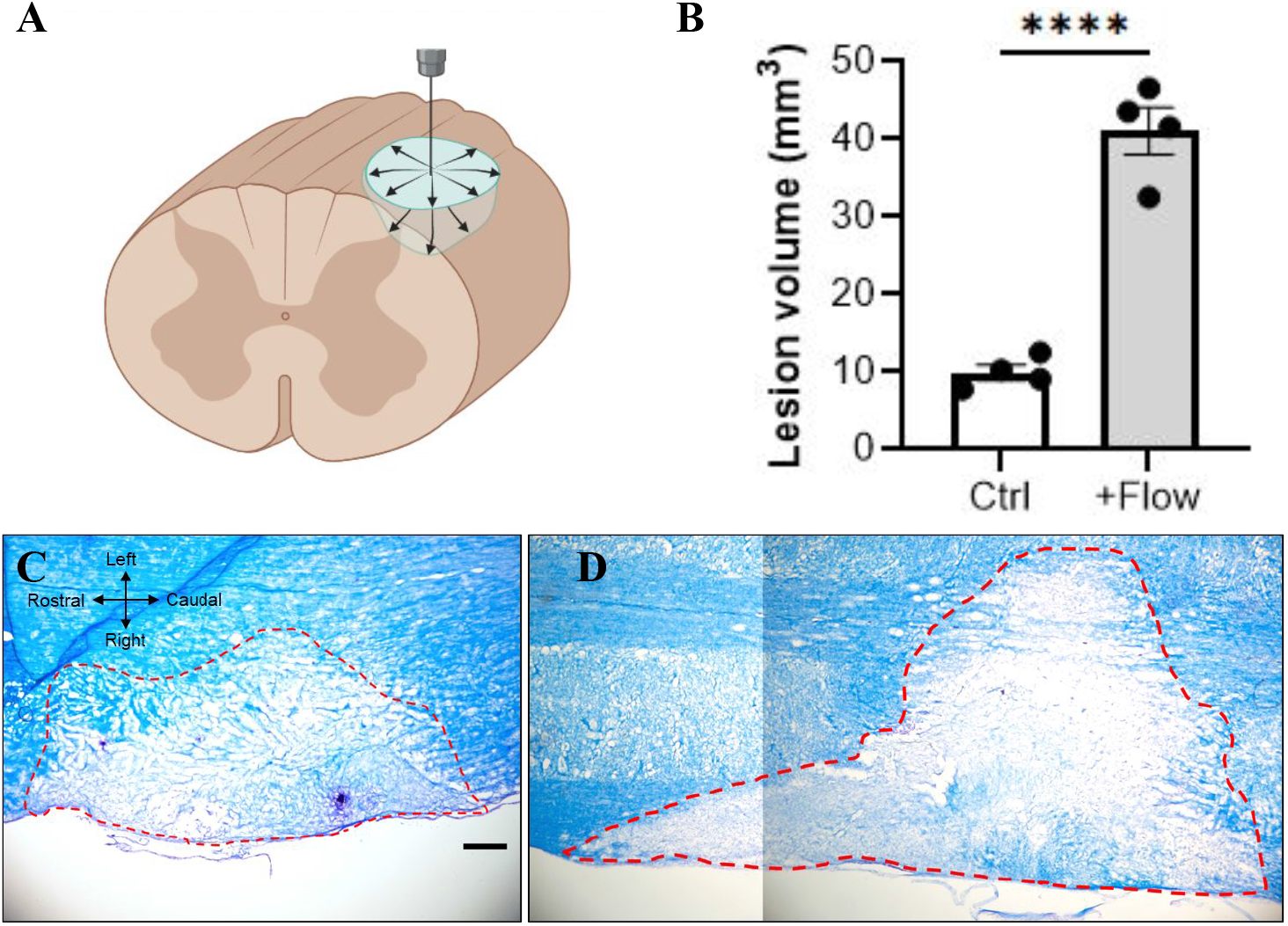
Adding convective interstitial flow at 7 DPI significantly increases lesion volume. A) Diagram showing the catheter-based technique to increase interstitial flow. Created using an institutional license for Biorender. B) Stereological quantification of spinal cord lesion volume with and without added convective flow.. C-D) Representative brightfield images of luxol fast blue stained tissue for the non-flow control group (C) and convective flow group (D). Scale bar is 200 μm. Groups were compared using unpaired t-tests. ****p<0.0001.

**Figure 6:**
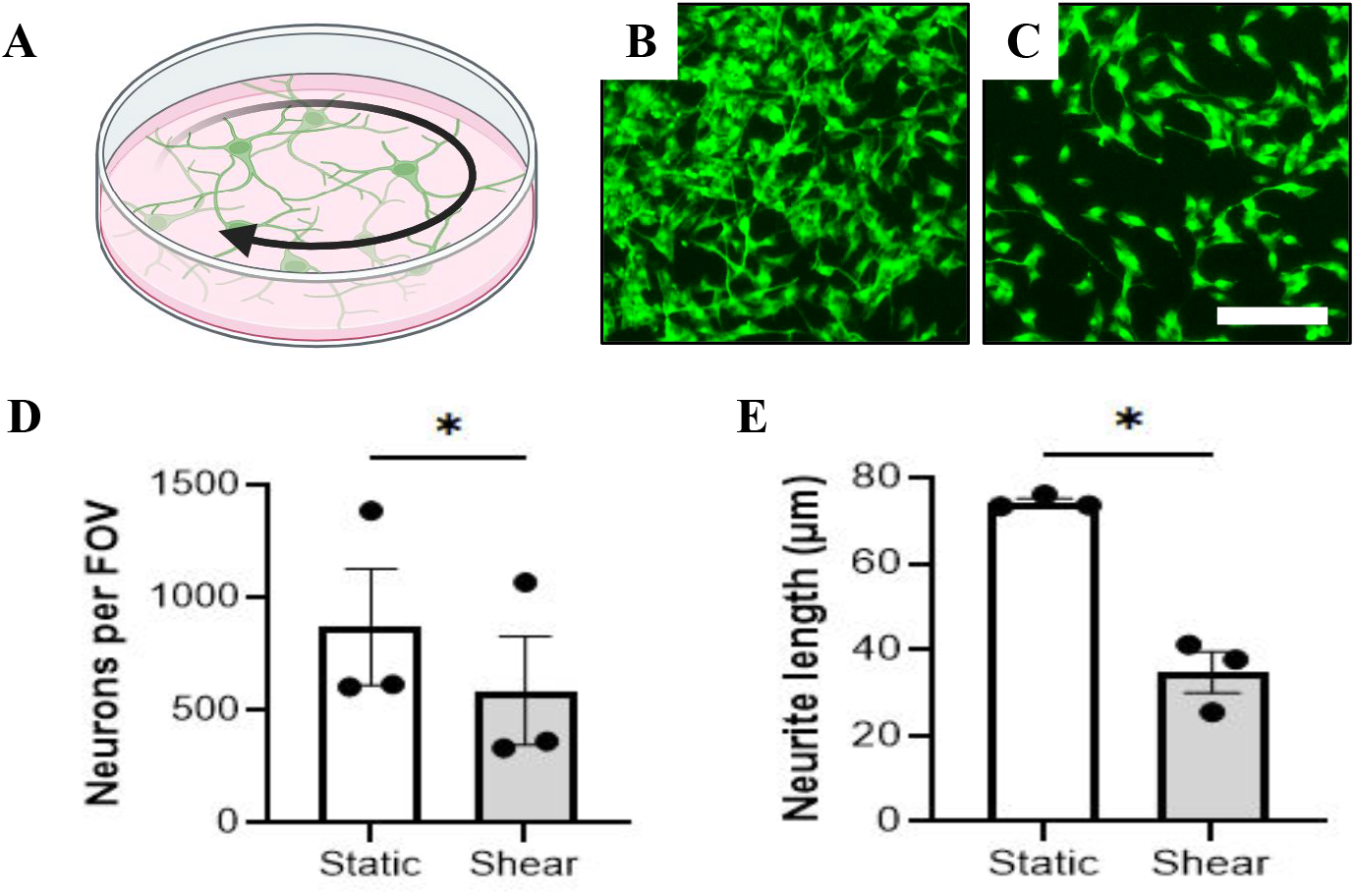
Exposure of neurons to low pathological fluid shear stress decreases cell count and neurite length. A) Diagram of neuronal culture under rotational fluid shear stress on an orbital shaker. Graphic made using an institutional license for Biorender. B-C) Representative fluorescence image of neurons cultured in static (B) and 0.1 dynes/cm^2^ shear (C) conditions overnight. Scale bar is 100 μm. D) Neuronal cell count per field of view (FOV). E) Neurite length traced using NeuronJ. Groups were compared using ratio paired t-tests. *p<0.05.

### Exposing neuronal cultures to pathological fluid shear decreases cell count and neurite length

Finally, we used *in vitro* cell cultures to validate the *in vivo* CED results and further establish the relationship between increased fluid flow and neural cell damage (**Figure 7**). We exposed differentiated SH-SY5Y neuronal-like cells overnight to a constant rotational shear stress of 0.1 dynes/cm^2^, the lower end of the pathological range of shear stresses predicted by our experiments and others.^22,32–34^ Representative images from the static and shear conditions are shown in **Figure 7B** and **C**. We find neuronal cell count is significantly reduced after overnight exposure to 0.1 dynes/cm^2^ of shear stress compared to the static controls (**Figure 7D**) (p<0.05). Furthermore, the average neurite length reduces from 75 μm in static to 35 μm with shear (**Figure 7E**) (p<0.05). Hence, shear stress either induces neurite retraction or prevents additional neurite extension. These results demonstrate that even a low pathological rate of fluid shear stress can influence neural cell survival and function.

## Discussion

Understanding the factors contributing to secondary injury can offer a therapeutic opportunity for protecting neural tissue and minimizing functional losses after SCI. Others have reported immediate and dramatic increases in intrathecal or intraspinal fluid pressure in rats, dogs, and humans.^9–11^ The water content starts to decrease after three days, and a significant, though perhaps not complete, relief is observed by seven days.^9,35^ This timeline coincides with lesion expansion or cavitation due to secondary injury: The largest tissue volume losses occur between 3 and 7 days, though the cavity can continue to expand for up to 14 days.^7,36^ An open question is if the pressure difference between injured and uninjured tissue will affect fluid flow within the tissue and whether such flow can contribute to secondary neural injury.

Intrathecal pressure is often used to approximate pressure within the spinal cord interstitium, and we tested the validity of this approach by measuring interstitial pressure (ISP) in the normal and injured rat spinal cord. We recorded an average ISP of -4 mmHg in the uninjured cord, compared to prior measurements at 2.7 mmHg in the rat intrathecal space.^25^ Negative values for interstitial pressure have been reported in several peripheral tissues in rats^37^ and would rationally facilitate transport of fluid and solute out of blood vessels and into the tissue. After injury, the interstitial pressure increased within one hour, peaked at 14 mmHg by 3 days, and returned close to baseline by 7 days. Khaing, et al., [2017] showed intrathecal pressure increases after injury to a peak of 8.9 mmHg for at least 3 days and levels off by 7 days.^25^ Interstitial pressure therefore appears to be higher than intrathecal pressure after injury, and so is the pressure differential that develops between injured and uninjured tissue. Nonetheless, this needs to be confirmed in humans, since Werndle et al., reported intrathecal pressure in humans to be over 20 mmHg at 24 hours post-injury.^38^

To understand the relationship between fluid pressure and interstitial fluid flow, we developed a computational fluid dynamics model of the rat spinal cord after injury. As one might expect, the dramatic increase in interstitial pressure by day 3 is predicted to cause an increase in interstitial flow velocity up to a maximum of 0.45 μm/s, or nearly 4.5 times faster than physiological interstitial velocity.^39^ This effect was significantly reduced in the 7 DPI model due to the lower pressure in the cavity. To somewhat validate these results and visualize interstitial flow *in vivo*, we exploited damage to the blood-spinal cord barrier to deliver the dye Evans Blue. This approach has been used repeatedly in the literature to visualize and demarcate regions of extravascular and interstitial flow *in vivo*.^40^ We find that Evans Blue infiltrates approximately four-fold more into the cord at 3 DPI than at 7 DPI, as measured perpendicular from the lesion boundary. These data agree with our COMSOL results, where fluid velocity was ∼3 times faster at 3 DPI than at 7 DPI. We are cautious, however, to use Evans Blue measurements to estimate a fluid velocity, since the two-dimensional measurement cannot account for the three-dimensional flow within real tissues. Additional studies are therefore needed using intravital imaging to more accurately quantify interstitial flow velocities *in situ* after SCI.

We hypothesized that increased flow will increase fluid shear stress within the tissue, which may contribute to cell damage after SCI. In the COMSOL model, the maximum velocity corresponds to a peak shear stress of ∼60 mPa or 0.6 dynes/cm^2^. This value is 6 times higher than physiological estimates and in a similar range predicted for tumors (0.1-1 dynes/cm^2^).^30,31,39^ Even at the lower end of this range, studies have shown that astrocytes upregulate inflammatory cytokines and microglia adopt pro-inflammatory reactive states.^41–43^ By day 7 after injury, the residual positive pressure is still predicted to induce fluid shear stresses twice that of physiological, indicating that neural cells surrounding a lesion may experience significant fluid forces for the duration of the subacute injury phase. We then evaluated spinal cord tissue for expression of cleaved caspase-3, a marker of apoptosis due to secondary injury. Both 3 and 7

DPI tissue showed robust expression of cleaved caspase-3, and this expression was moderately to strongly colocalized with fluorescence of Evans Blue. Therefore, regions of cell death correlate with regions of elevated interstitial flow during subacute secondary injury.

To assess a potential causative role for elevated interstitial flow and cell damage, we artificially increased fluid flow in the lesion using an exogenous infusion at 7 DPI, after interstitial pressure approximately normalizes.This technique of convection enhanced delivery, or CED, is used clinically to enhance drug perfusion in high-pressure brain tumors and has been shown experimentally to increase flow-stimulated cellular signaling.^32^ Applying CED after SCI at three times a physiological flow rate (1 μL/min) caused a 4-fold increase in lesion volume. This result signifies that high interstitial flow contributes to or exacerbates tissue damage. While the complete mechanism is yet unclear, our *in vitro* experiments demonstrate that exposing neurons to low pathological shear stress (0.1 dynes/cm^2^) results in shorter neurite extensions and reduced neuronal cell count compared to static controls. Future work will elucidate if the pathological effects of fluid flow and shear stress on neural cells and tissue is due to mechanical effects, transport effects, or both. Furthermore, our results suggest that direct fluid infusion (e.g., drug or cell administration) into an SCI cavity should be minimized to reduce the risk of tissue damage.

## Materials and Methods

### Animal Procedures

All animal procedures were approved and conducted in accordance with the guidelines of the Institutional Animal Care and Use Committee of the University of Massachusetts Amherst, as well as in compliance with the National Institute of Health’s Guide for the Care and Use of Laboratory Animals. Sprague-Dawley rats (8–12 weeks of age) of both sexes were purchased from Charles River Laboratories. A cervical C4 hemi-contusion spinal cord injury was performed to simulate the most common spinal cord injury observed in the clinic.^44^ Rats were induced under anesthesia with 3% isoflurane and maintained under 1-2% isoflurane. The cervical area was shaved and aseptically prepared for surgery using alternating washes of chlorhexidine and sterile saline. The rats were placed on a diaper pad throughout the surgical procedure. A cervical laminectomy of C3-C5 was performed aseptically, followed by a C4 hemi-contusion at 150 kDynes using an Infinite Horizons spinal impactor (IH-0415, World Precision Instruments). Sham animals underwent all procedures with the exception of the spinal impact. The overlying muscle and skin were then sutured, and the rats were allowed to recover in their home cage. The rats received analgesic (0.1 mg/kg, buprenorphine, Par Pharmaceutical) subcutaneously for 72 hours after the operation and were given dietary support (Dietgel) and sterile saline for hydration, as needed. At experimental endpoints, rats were euthanized using carbon dioxide asphyxiation to avoid artifacts in flow due to transcardial perfusion.

To measure intraspinal pressure, the dura and spinal cord was punctured using a 20G needle, and a pressure measuring catheter (3.5F, Single, Straight, 100cm, ADinstrument®) was inserted. The data were recorded with software PowerLab® (ADinstrument®) for 30 minutes each time. The probe was calibrated to read 0.5 mV as 0 mmHg and 3.0 mV as 100 mmHg. In separate animals, the pressure was measured in the naive spinal cord (n=3), 1 hour post-injury (HPI, n=5), 3 days post-injury (DPI, n=5), and finally 7 DPI (n=3).

In one cohort, Evans Blue (Sigma-Aldrich) was administered at 2 mg/kg via tail vein injection 24 hours prior to tissue harvest to label regions vascular leakage and interstitial flow *in vivo*. In another cohort, interstitial flow was artificially enhanced at 7 DPI, after fluid pressure approximately normalized, using convection enhanced delivery (CED). Briefly, the rats received the same C4 hemi-contusion injury described above and were allowed to recover for 7 days. The control group had their injury reopened but only received a cavity puncture using a 25G needle. The +Flow group received reopening of injury, puncturing of the cavity, and delivery of sterile saline at 1 µL/min (approximately three times the normal interstitial flow rate).^45,46^ Both groups were re-sutured and kept in their home cage for an additional 3 days. Ketoprofen (1 mg/kg, Zoetic©) was administered subcutaneously for 48 hours after the CED. The spinal cords were harvested at 72 hours after CED. For all studies, the spinal columns were harvested and fixed in 4% paraformaldehyde for 24 hours and cryopreserved in 30% sucrose until sectioning.

### Histology and Immunohistochemistry (IHC)

Fixed spinal cords were frozen in Optimum Cutting Temperature medium and cryosectioned longitudinally at 20 µm thickness using a Leica CM1950. For IHC, tissues were permeabilized using blocking buffer (0.3% Triton X-100, 3% donkey serum in PBS) for 1 h at room temperature. The slides were then labeled overnight at 4℃ with primary antibodies for neurofilament-heavy (RT-97, Developmental Studies Hybridoma Bank, University of Iowa) or cleaved caspase-3 (Cell Signaling 9664S). After three rounds of washing, secondary antibodies were applied with the blocking buffer in the dark for 1 hour. Sections were again washed three times in 1x phosphate buffered saline (PBS), counterstained with 4’,6-diamidino-2-phenylindole (DAPI, 1:10,000) for 5 minutes, and mounted using Fluoromount-G (Southern Biotech). Images were acquired using a Nikon A1R confocal microscope with 10x or 20x air objectives. Four to six slices were imaged and analyzed, one image per slice.

CED tissues were stained using Luxol Fast Blue. Briefly, the slides were incubated in 0.1% Luxol Fast Blue solution (Thermo Scientific) at 56℃ for 14 hours. Excess stain was rinsed using 95% ethyl alcohol and deionized water, and the stain was differentiated using 0.05% lithium carbonate solution (Thermo-Fisher) for 30 seconds followed by 70% ethyl alcohol. The slides were then washed with deionized water and counterstained with 0.1% Cresyl echt violet solution (Thermo-Fisher) for 50 seconds. After the final staining, tissues were dehydrated by washing once for 5 minutes with 95% ethyl alcohol, twice with 100% ethyl alcohol, and finally twice in 100% xylene. The stained slides were viewed and imaged with a compound light microscope (Leica Microsystems). The lesion volume was estimated using stereology and ImageJ software. A total of 24 sections were measured in each rat to estimate the volume of the lesion.

### COMSOL Modeling

COMSOL Multiphysics and the porous media module were purchased under an individual license. The software was used to develop a computational model of the rat spinal cord to assess fluid velocity and shear stresses in the context of spinal cord injury. While not an exhaustive representation, the *in silico* model enabled simulating how fluid pressure at different time points affects interstitial fluid flow out of an idealized lesion cavity and calculating the resulting fluid velocities and shear stresses. The cord parenchyma was modeled as a cylinder of 3 mm radius and a length of 60 mm to minimize end effects. We chose a simplified model of the white matter, which may degenerate more than gray matter in cervical enlargements.^47^ The lesion was modeled as a sphere with diameter 1.5 mm, approximating prior lesion volume measurements of 1.77 μL, set in the upper right quadrant to mimic the *in vivo* injury paradigm. The pressure measurements from the above experiment and parameters from the literature (**Table 1**) were entered into the model to define the flow parameters. Different runs with distinct pressures of different time points of SCI were conducted. The spinal pressure away from the injury at 3 DPI and 7 DPI were set as normal spinal pressure. The computations were performed using the steady-state Brinkman’s equation model:

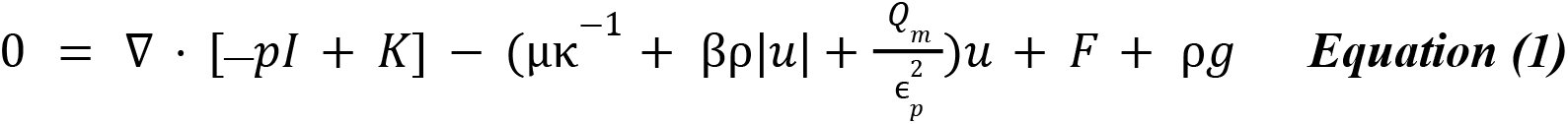

Shear stress in the tissue was calculated by the method presented by Guyot, et al. [2015].^48^ The porous tissue is approximated as a composite of cylindrical ducts with diameter δ, such that the velocity profile is similar to Poiseuille flow. Additionally, superficial velocity is divided by tissue porosity (ε) to obtain a discrete velocity (*u*). Shear stress is then be calculated as follows:

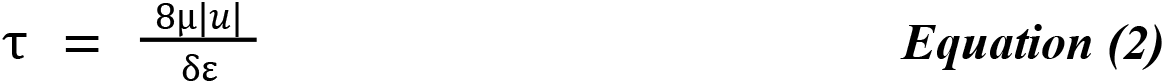

The graph was acquired by calculating the shear stress values across a line drawn through the epicenter of the cavity in the longitudinal direction.

### Image Analysis

The distance traveled by extravasated Evans Blue dye beyond the lesion margin was measured as a proxy for the path and rate of interstitial flow after injury. The lesion area was demarcated based on a lack of signal intensity in DAPI and/or neurofilament staining. The distance was measured using ImageJ by drawing a perpendicular line from the edge of the lesion area. The distances were averaged for each slice and plotted such that each dot represents an individual animal.

Nikon NIS-elements software was used for colocalization analysis. A region of interest was drawn to exclude effects of staining or imaging artifacts, and Pearson’s correlation coefficient (R_p_) and Mander’s colocalization coefficients (k_1_, k_2_) were calculated for each image. The most commonly used quantitative estimate of colocalization, Pearson’s correlation coefficient (R_p_), depicts the degree of overlap between two selected channels. A positive number for the R_p_ represents the degree of overlap of fluorescent signals, with 1.0 representing complete overlap and zero representing random placement or no overlap. Mander’s colocalization coefficients define the interacting fraction for that image. Three images at 100 μm apart were analyzed and averaged per animal.

### Cell Culture and Immunocytochemistry

Human neuroblastoma SH-SY5Y cells were purchased from ATCC (Manassas, VA) and differentiated into neuronal-like cells for *in vitro* shear experiments. The cells were cultured using a maintenance medium of 1:1 Dulbecco’s Modified Eagle Medium: F12 nutrient supplement with 10% fetal bovine serum and 1% non-essential amino acids. At the time of experiment, 50,000 undifferentiated cells were plated into wells of two 6-well plates. Both plates were treated for five days with differentiation medium consisting of Neurobasal medium plus 0.25% Glutamax, 1% B27, 0.5% N2, and 10 μM of freshly diluted all-trans retinoic acid (Fisher Scientific). The medium was refreshed at least once during the differentiation period. On day five, the cells in one plate were exposed to 0.1 dynes/cm^2^ shear stress for 18 hours using an orbital shaker at 100 rpm.^49^ The other plate was kept in static condition as a control. Both plates were then fixed using 4% paraformaldehyde for 20 minutes at room temperature and stored in 1X PBS until ready for staining. The samples were treated for 1 hour at room temperature with a blocking buffer of 0.3% Triton X-100 and 3% donkey serum in 1X PBS. The primary anti-beta III tubulin antibody (Abcam ab7751) was added into the blocking buffer, and the samples were incubated overnight at 4℃. The samples were then washed three times with 1X PBS for ten minutes each and incubated in the dark for one hour at room temperature with a donkey anti-mouse secondary (ThermoFisher A-21202). The cells were washed with 1X PBS as before, counterstained with DAPI, and stored in 1X PBS. Each well was imaged in four random but non-overlapping locations around the outer edge (peak shear stress). Cell count per field of view (FOV) and neurite length were measured using ImageJ. Experiments were conducted with three technical replicates for three biological replicates each.

### Statistical Analyses

All data were graphed and analyzed using GraphPad Prism 10 software. The modes of intraspinal pressure measurements were analyzed by one-way analysis of variance (ANOVA) and multiple comparisons with Tukey’s correction. All other *in vivo* data were compared using student t-tests. *In vitro* groups were compared using paired t-tests. For all graphs, error bars represent the standard error of the mean (SEM). Each data point represents a biological replicate (number of rats or number of cell culture replicates). Statistical significance is indicated by *p < 0.05, **p < 0.01, ***p < 0.001, ****p < 0.0001.

## Conclusions

Traumatic injury to the central nervous system induces neural tissue degeneration, which leads to disability and loss of function.^3–5^ In the acute phase of spinal cord injury, influx of water content to the lesion results in the formation of edema above the cord.^9,10^ In this study, we showed that fluid pressure is also increased within the interstitial region after injury. Using simulations, we predict that the increase in interstitial fluid pressure causes increased interstitial fluid flow, particularly 3 days after injury. These results were validated *in vivo* based on quantifying a proxy marker of vascular leakage. Regions of interstitial flow after injury correlated with the apoptosis marker cleaved caspase-3, suggesting that secondary cell damage is related to fluid flow in the cord. We used an *in vivo* flow manipulation technique to demonstrate that an increase in flow increases tissue damage, and an *in vitro* model that pathological fluid shear stress decreases neuronal cell survival and neurite length. Future research will help define the mechanisms by which fluid flow and shear stress mediate tissue damage. Nonetheless, our results provide new insight into the basic pathophysiology of SCI and may inform therapeutic development to protect neural tissue from secondary injury.

## Authorship Contributions

RCC conceptualized the project idea; HYK and RCC developed the investigational plan and methodology; HYK and CS conducted the experiments for data acquisition and analysis; RCC supervised experimental design and data analysis; HYK and RCC developed the original manuscript draft and curated data for publication; RCC acquired funding and managed the procurement of reagents and supplies; all authors edited and reviewed the final manuscript.

## Acknowledgements

We would like to thank James Chambers and the UMass Nikon Center of Excellence for microscopy assistance; Dr. Paul Spurlock, Joanne Huyler, Cathy Cervi, and the rest of the vet care staff at UMass for their assistance and guidance with animal care and surgical procedures; Mary Chase Sheehan and Dr. Govind Srimathveeravalli for advice in COMSOL modeling.

## Data Sharing And Data Availability

Data supporting the *in vivo* findings of this study are openly available in the Open Data Commons for Spinal Cord Injury. All other data are available upon request to the corresponding author.

