## Supplemental Figure 1 for "Convective forces contribute to post-traumatic degeneration after spinal cord injury"

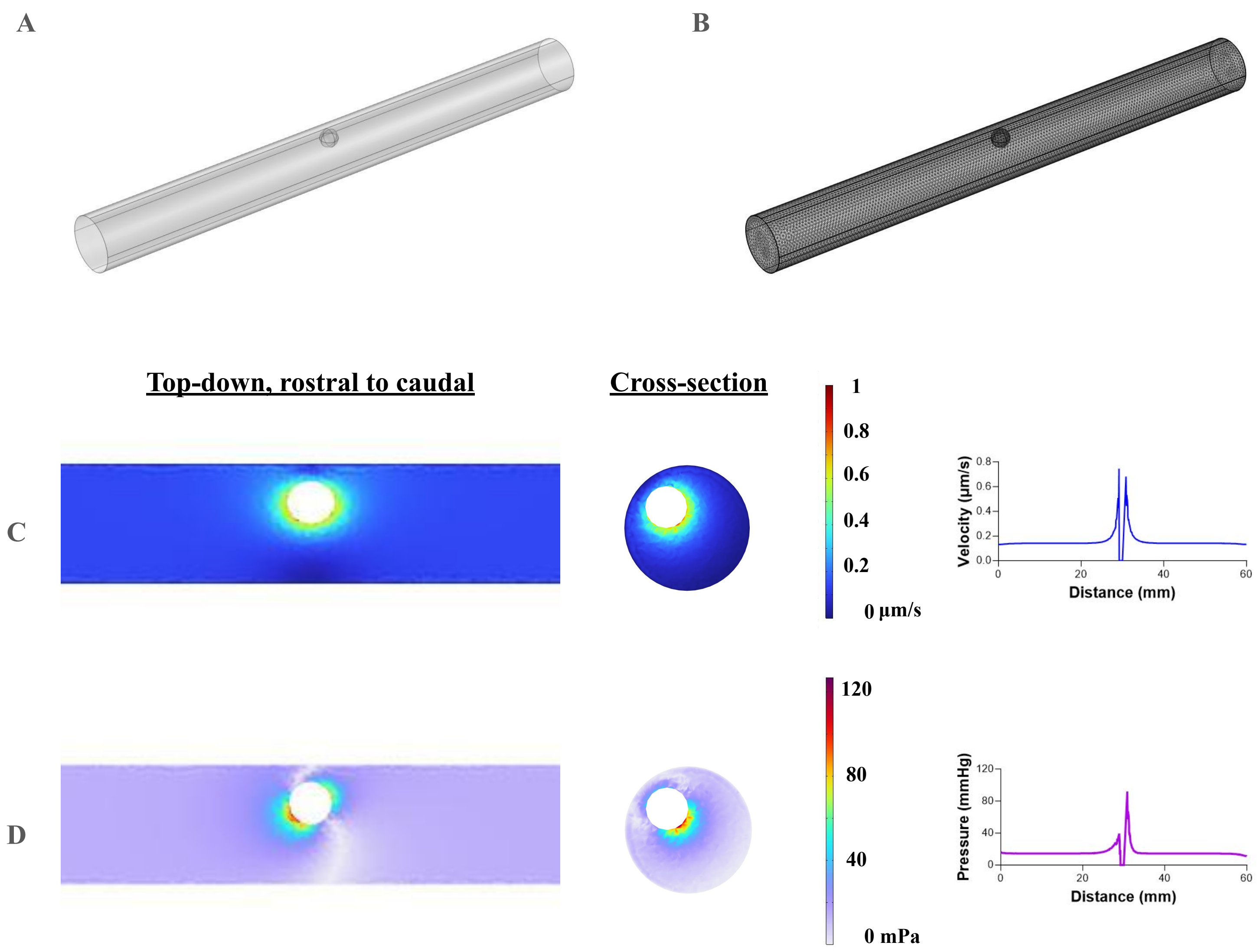


**Supplemental Figure 1.** A) Representation of the COMSOL model. B) Mesh generated with finer setting. 593886 domain elements, 21320 boundary elements, and 824 edge elements. (C-D) Velocity (C) and shear stress (D) of 3 DPI model without incoming interstitial fluid flow in the +x direction. The peak magnitude is higher compared to the IFF model, and the flow is isotropic in the x direction.
